# Bioenergetic profiling of fresh human kidney tissue reveals compensatory metabolic adaptation and intrinsic mitochondrial dysfunction in diabetes

**DOI:** 10.64898/2026.07.23.740003

**Authors:** Cesare Granata, Adrienne Laskowski, Vicki Thallas-Bonke, Georg Ramm, Richard J. MacIsaac, Christopher Chang, Nicholas Campbell, Peter Royce, Mark E. Cooper, Elif I. Ekinci, Jeremy Grummet, Scott G. Wilson, Catriona A. McLean, Melinda T. Coughlan

**Affiliations:** Department of Diabetes, School of Translational Medicine, Monash University, Alfred Research Alliance, Melbourne, Victoria, Australia; Membrane Biology Group, Department of Biochemistry and Molecular Biology, Monash University, Clayton, Victoria, Australia; Department of Endocrinology & Diabetes, St Vincent’s Hospital Melbourne, Fitzroy, Victoria, Australia; Department of Medicine, St Vincent’s Hospital Melbourne, University of Melbourne, Fitzroy, Melbourne, VIC 3065, Australia; Australian Centre for Accelerating Diabetes Innovations, School of Medicine, University of Melbourne, Parkville, Melbourne, VIC 3052, Australia; Department of Urology, Alfred Health, Melbourne, Victoria, Australia; Department of Surgery, School of Translational Medicine, Monash University, Alfred Research Alliance, Melbourne, Victoria, Australia; Department of Medicine, The University of Melbourne, Parkville, Victoria, Australia; Department of Endocrinology, Austin Health, Heidelberg, Victoria, Australia; Department of Medicine, School of Translational Medicine, Monash University, Alfred Research Alliance, Melbourne, Victoria, Australia; Department of Renal Medicine, Alfred Health, Melbourne, Victoria, Australia.; Department of Anatomical Pathology, Alfred Health, Melbourne, Victoria, Australia.; Baker Heart & Diabetes Institute, Melbourne, Victoria, Australia; Drug Discovery Biology, Monash Institute of Pharmaceutical Sciences, Monash University Parkville Campus, Parkville, Victoria, Australia

## Abstract

The kidney is a highly energetic organ, requiring substantial ATP production through mitochondrial oxidative phosphorylation to support tubular reabsorption. Metabolic reprogramming and impaired mitochondrial function are implicated in diabetic kidney disease, yet direct assessment of mitochondrial respiratory flux in the human kidney has been constrained by limited access to freshly obtained tissue. Consequently, much of the evidence supporting altered renal mitochondrial function in diabetes derives from animal models that do not fully recapitulate the human condition. We established a workflow for real-time bioenergetic profiling of fresh kidney cortex obtained during nephrectomy from living individuals with diabetes and preserved kidney function. Mitochondrial respiration, electron transport system activity and tubular mitochondrial morphology were compared with age- and sex-matched, histopathologically normal non-diabetic controls. High-resolution respirometry revealed increased mitochondrial respiratory flux in permeabilised diabetic kidney cortex. In contrast, mitochondria isolated from the same tissue exhibited reduced respiratory capacity and impaired complex I activity. Quantitative analysis of tubular cells demonstrated increased mitochondrial volume density together with greater mitochondrial fragmentation in diabetes. These findings reveal that the human kidney undergoes substantial metabolic adaptation early in diabetes, before measurable loss of kidney function. Increased tissue-level respiratory flux despite intrinsic mitochondrial impairment suggests that expansion and remodelling of the mitochondrial network may initially compensate for reduced organelle efficiency and sustain the kidney’s high energetic demands. This compensatory state may, however, increase metabolic stress and vulnerability to subsequent kidney injury. To our knowledge, this study provides the first direct tissue-level functional evidence that mitochondrial metabolism is reprogrammed in the human kidney in diabetes before measurable kidney dysfunction develops. It defines an early bioenergetic signature characterised by tissue hypermetabolism despite impaired mitochondria-specific respiratory capacity, challenging the concept that diabetes produces a uniform decline in renal mitochondrial function. Failure to sustain this adaptive state may represent a critical transition towards diabetic kidney disease.

**GRAPHICAL ABSTRACT:** 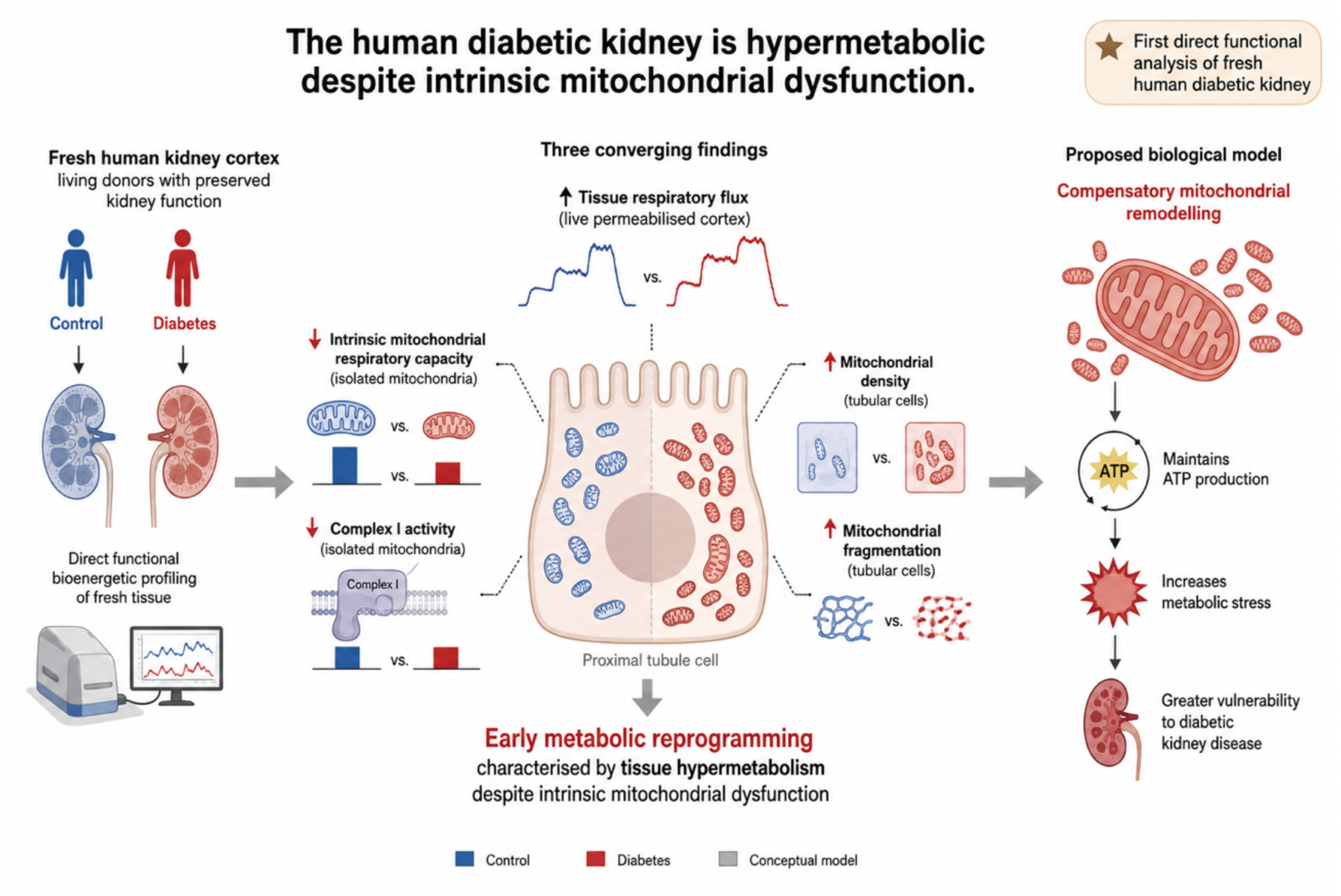

**One Sentence Summary:** Diabetes drives early metabolic reprogramming of the human kidney before measurable kidney dysfunction

## Introduction

Currently, one in ten adults is affected by diabetes, and its global prevalence continues to increase ^1^. Diabetic kidney disease (DKD) affects more than 30% of people with diabetes and is the most common cause of end stage kidney disease (ESKD) worldwide^2^. Despite recent advances in the clinical management of kidney disease ^3,4^ post-diagnosis, a significant proportion of patients still progress to ESKD and require renal replacement therapy, imposing a major health and economic burden.

Mitochondria are fundamental to metabolic homeostasis by converting nutrient-derived substrates into adenosine triphosphate (ATP) through oxidative phosphorylation (OXPHOS). Mitochondrial health is particularly important in the kidney, which relies heavily on OXPHOS to generate the ATP required for tubular reabsorption of filtered solutes ^5^. Indeed, the kidney consumes a disproportionately large amount of the body’s energy to support active sodium transport across the proximal tubular epithelium ^5^. Accordingly, poor mitochondrial health has been linked to kidney disease ^6^.

There is increasing evidence to indicate that the disruption in mitochondrial respiratory flux and energy generation (bioenergetics) is key to the development and progression of DKD ^7–17^. Experimental studies indicate that renal mitochondrial respiratory function declines before overt structural kidney injury develops, with reported abnormalities including reduced electron transport system (ETS) complex I activity, altered mitochondrial reactive oxygen species generation and ATP depletion ^12,13,15^. In human DKD, mitochondrial perturbations have been inferred from urinary metabolomic signatures and altered renal expression of mitochondrial enzymes and metabolic pathways ^7,18–21^. Recent advances in functional imaging and single-cell transcriptomic analyses have further revealed early alterations in kidney oxidative metabolism and tubular metabolic signalling in people with diabetes. However, these approaches provide indirect, molecular or whole-organ assessments and do not directly quantify mitochondrial respiratory flux in viable human diabetic kidney tissue. Consequently, current understanding of renal mitochondrial bioenergetics in diabetes continues to derive predominantly from experimental models.

Human kidney studies are further constrained by reliance on tissue obtained from deceased donors or archived biopsies from patients with established kidney disease. Such samples provide limited insight into the early metabolic adaptations that precede measurable loss of kidney function. Because mitochondrial respiration is highly dynamic and rapidly altered by tissue handling and preservation, fresh viable tissue is essential for direct assessment of bioenergetic function.

Here, we established a workflow for the rapid collection and real-time bioenergetic profiling of fresh kidney cortex obtained from living donors with type 2 diabetes and preserved kidney function. We directly quantified respiratory flux in fresh permeabilised kidney cortex and compared these findings with intrinsic respiratory capacity and electron transport system activity in mitochondria isolated from the same tissue.

Diabetic donors were compared with age- and sex-matched, histopathologically normal non-diabetic donor controls. By integrating tissue-level respiration, isolated mitochondrial function and quantitative analysis of tubular mitochondrial morphology, we identified an early bioenergetic phenotype characterised by increased tissue respiratory flux despite impaired intrinsic mitochondrial capacity, together with increased mitochondrial density and fragmentation. These findings provide direct functional evidence that the human kidney undergoes substantial mitochondrial and metabolic reprogramming in diabetes before measurable kidney dysfunction develops. These studies provide novel translational insight into oxidative phosphorylation and respiratory chain function in the human diabetic kidney, revealing early tissue hypermetabolism despite intrinsic mitochondrial dysfunction.

## METHODS

### Study Approval

This study was performed in accordance with the principles of the Declaration of Helsinki and was approved by the Ethical Review Committee of Alfred Health (Project number 474/16), Monash University (Project number 7760) and the Victorian Cancer Biobank (Project number 17014), Melbourne, Australia. Written informed consent was obtained from participants prior to inclusion in this study.

### Study Design

This was a cross-sectional observational study performed in Melbourne, Australia, involving individuals with normal glucose tolerance and subjects with diabetes.

### Patient Recruitment

From March 2017 to March 2020, patient recruitment took place at five hospitals across metropolitan Melbourne, Australia: 1) Alfred Health, 2) Eastern Health, 3) Peter MacCallum Cancer Centre, 4) Austin Health, and 5) Monash Health, following nonemergent radical or partial nephrectomy due to renal cell carcinoma, in an effort coordinated by the research group at Monash University, Alfred Health and the team at the Victorian Cancer Biobank (Melbourne, Victoria, Australia). Prospective patients were approached in the perioperative clinic for inclusion in the study if they met the following inclusion criteria: 1) Scheduled for impending surgery for the excision of renal cell carcinoma and, 2) aged between 18 to 80. Patients underwent a process of full informed consent during their perioperative consultation. The study was approved by the Ethical Review Committee of Alfred Health (Project number 474/16), Monash University (Project number 7760) and the Victorian Cancer Biobank (Project number 17014), Melbourne, Australia.

### Risk of bias assessment

No blinding method was included in the original design for this observational study. However, data and sample collection were carried out in and analysis was done in Monash University by separate scientists. The scientist evaluating the samples for Complex I and citrate synthase activity was blinded to the clinical parameters at the time of analysis.

### Exclusion criteria

Individuals outside the inclusion age range (18-80 years), (2) patients undergoing emergency (nonelective) surgery, (3) patients diagnosed as having a renal carcinoma with evidence of local invasive disease or unclear radiological margins precluding macroscopic identification of uninvolved renal cortical tissue, (4) suspected or confirmed nontumorous infiltrative renal disease.

### Biospecimen Collection

During nephrectomy, the kidney was collected by the Urological surgeon and immediately transported on ice to Anatomical Pathology. After oversight and handling of the specimen, the Pathologist selected the macroscopically uninvolved renal parenchyma, which was placed on ice and transported to the Monash University Diabetes department laboratory. Upon receipt, the kidney biospecimen was divided in the following portions: 1) ∼10-15 mg, placed in respiration buffer (MiRO5, see below) for determination of mitochondrial respiration in permeabilized tissue, 2) ∼2 mg was cut into ∼1 mm^3^ pieces and immersed in fixative for transmission electron microscopy (TEM), 3) ∼20-30 mg was placed in mitochondrial isolation buffer (see below) for determination of mitochondrial respiration in isolated mitochondria, and 4) the remaining sample was cut in small 10-20 mg pieces and immediately frozen in liquid nitrogen for subsequent enzyme activity. The time lapse between obtaining the tissue from pathology to sample dissection for fresh or frozen analyses was between 1 and 3 h.

### Transmission electron microscopy for analysis of mitochondrial morphology

Mitochondrial morphology was determined in electron micrographs of proximal tubule epithelial cells. Immediately upon receipt of human kidney cortex from pathology, 4-6 pieces of 1 mm^3^ renal cortex were fixed in 1 mL of electron microscopy (EM) primary fixative buffer (0.1 M cacodylate buffer pH 7.4, 2% paraformaldehyde, 2.5% glutaraldehyde, 4.5 mM calcium chloride) for 1 h at 25 °C, followed by 24 h at 4 °C in 1 mL of fresh EM primary fixative buffer. The next day samples were washed 3 times with 0.2 M cacodylate buffer pH 7.4, and sent to the EM facility for secondary fixation, which was performed using 1% osmium tetroxide and 1.5% potassium ferricyanide in 0.1 M cacodylate buffer for 1 h at 25 °C. Samples were finally rinsed in three washes of MilliQ H_2_O for 15 minutes each.

The fixed tissues were dehydrated by incubating in increasing concentrations of ethanol for 15 min, consisting of 30, 50, 70, 90 and 100% ethanol. The ethanol was removed and replaced with 100% propylene oxide. Dehydrated tissues were incubated in a mixture of Epon resin and propylene oxide at a ratio of 1:1 for 6 h at room temperature, followed by a 2:1 Epon/propylene oxide mixture overnight. Tissues were incubated in 100% freshly made Epon resin for 6 h, followed by another 100% resin change overnight. The tissues were then placed into Beem capsules in 100% resin and the resin polymerized for 48 h in an oven at 60 °C.

Resin embedded tissue was sectioned with a Diatome diamond knife using a Leica UCS ultramicrotome. Sections of thickness 70 – 90 nm were collected onto formvar-coated 100 mesh copper grids and stained sequentially with 1% uranyl acetate for 10 minutes and lead citrate for 5 min. Sections were imaged in a JEOL 1400+ transmission electron microscope at 80 kV, and images captured with a digital camera at a resolution of 2K x 2K. The image field size was 17.6 × 17.6 μm. Individual mitochondria were circled by hand using the (Fiji Is Just) ImageJ Version 1.49m software (National Institutes of Health, Bethesda, MD, USA) from n = 5 images/patient, for a total of 18 Ctrl and 14 T2D patients. Several shape descriptors were assessed: surface area (in µm^2^); perimeter (µm); circularity, determined as [4π·(surface area/perimeter^2^)] and roundness, determined as [4·surface area/(π·major axis^2^)], both reflecting sphericity, with 1 indicating perfect spheroids; aspect ratio, determined as [(major axis)/(minor axis)] and reflecting the “length-to-width ratio”; Feret’s diameter (µm), which reflects the longest distance between any two points pertaining to a given mitochondrion; form factor, determined as [perimeter^2^/(4π·surface area)], reflecting mitochondria branching and complexity ^22^. In addition, Mito_VD_ was determined as the sum of all individual mitochondrial areas within a given image, normalized by the cellular area (i.e., the extracellular space area was removed) subtracted of the nuclear area. Mitochondria that were cut at the border of the image were also circled and were used for determination of Mito_VD_, but were excluded from all shape descriptor determination.

### Sample preparation for high-resolution respirometry in permeabilized kidney cortex

Fresh kidney cortex was mechanically separated at 4 °C in MiR05 (0.5mM EGTA, 3mM MgCl_2_, 60mM K-lactobionate, 20mM taurine, 10mM KH_2_PO_4_, 20mM HEPES, 110mM sucrose and 1 g/L BSA essentially fatty acid-free, pH 7.1), followed by permeabilization by gentle agitation for 30 min at 4 °C in MiR05 containing 100 μg/mL of saponin, and 3 5-min washes in MiR05.

### Mitochondrial respiration protocol

Mitochondrial respiration was determined in duplicate (from 2.5-5 mg wet weight of kidney cortex) in MiR05 at 37 °C using the high-resolution Oxygraph-2k (Oroboros, Innsbruck, Austria). To avoid potential oxygen diffusion limitation, oxygen concentration was maintained between 270 and 480 nmol mL^-1^ by re-oxygenation via direct syringe injection of O_2_. A substrate-uncoupler-inhibitor titration (SUIT) protocol was used, as previously described ^23^, with minor modifications: 0.2 mM octanoylcarnitine and 2 mM malate to stimulate leak respiration (L) in the absence of adenylates and limitation of flux by electron input through ETF ([ETF]_L_); 3 mM MgCl_2_ and 5 mM ADP to stimulate OXPHOS capacity (P) with saturating levels of ADP and limitation of flux by electron input through ETF ([ETF]_P_); 5 mM pyruvate and 10 mM glutamate to stimulate P with saturating levels of ADP and limitation of flux by convergent electron input through ETF + CI ([ETF+CI]_P_); 10 mM succinate to stimulate maximal P with saturating levels of ADP and limitation of flux by convergent electron input through ETF + CI + CII ([ETF+CI+II]_P_); 10 μM cytochrome *c* to test the outer mitochondrial membrane integrity (chambers with a >10% flux increase were excluded); 2.5 µM oligomycin (an inhibitor of CV) to stimulate L in the presence of adenylates with limitation of flux by convergent electron input through ETF + CI + CII ([ETF+CI+II]_L_); 1-2 μM carbonyl cyanide 4-(trifluoromethoxy) phenylhydrazone (FCCP) via stepwise titration to stimulate maximal electron transport chain capacity [E] with saturating levels of ADP and limitation of flux by convergent electron input through ETF + CI + CII ([ETF+C+II]_E_); 0.5 μM rotenone (an inhibitor of CI) to stimulate E with saturating levels of ADP and limitation of flux by electron input through CII ([CII]_E_); 5 μM antimycin A (an inhibitor of CIII) to determine residual non-mitochondrial oxygen consumption. Data are presented as mass-specific mitochondrial respiration (pmol O_2_ s^-1^ mg^-1^ wet weight) and as mitochondria specific respiration (pmol O_2_ s^-1^ mg^-1^ wet weight/Mito_VD_).

### Respiratory flux control ratios

To pinpoint the origin of the differences in mitochondrial respiration between groups the following FCRs were calculated: 1) the LCR was determined as the quotient of [ETF+CI+II]_L_ over [ETF+CI+II]_E_; 2) the PCR was determined as the quotient of [ETF+CI+II]_P_ over [ETF+CI+II]_E_; 3) the inv-RCR was determined as the quotient of [ETF]_L_ over [ETF]_P_; 4) a second inv-RCR was determined as the quotient of [ETF+CI+II]_L_ over [ETF+CI+II]_P_.

### Mitochondrial isolation for high-resolution respirometry and enzymatic activity determination

Mitochondrial isolation was performed on ∼20-30 mg of fresh (for determination of mitochondrial respiration) or ∼30-40 mg of frozen (for determination of enzymatic activity) kidney cortex. In both cases, samples were minced with a sharp blade at 4 °C in mitochondrial isolation buffer (70 mM sucrose, 210 mM mannitol, 5 mM HEPES, 1 mM EGTA, pH 7.2; for isolation from fresh samples for determination of mitochondrial respiration 0.5% (w/v) essentially fatty acid free bovine serum albumin (BSA) was added fresh to the above buffer, and subsequently lysed with 8-10 strokes in a Potter-Elvehjem tissue grinder. Following centrifugation for 10 min at 4 °C at 800 g the supernatant was collected and further centrifuged for 20 min at 4 °C at 8000 g. The supernatant was discarded and the pellet was washed in 500 mL of fresh isolation buffer and centrifuged a second time with the same settings. The ensuing pellet was resuspended in 150 mL of isolation buffer. To minimize downtime, isolated mitochondria obtained from fresh kidney cortices were immediately used for high-resolution respirometry and protein concentration was determined a posteriori (BCA Protein Assay Kit, Pierce-Thermo Fisher Scientific, Melbourne, Victoria, Australia).

Conversely, for isolated mitochondria fractions from frozen kidney cortices, protein concentration was determined immediately before determination of enzymatic activity.

### High-resolution respirometry in mitochondria isolated from kidney cortex

Mitochondrial respiration was measured in duplicate (from 10-30 µg of mitochondrial-enriched protein) in MiR05 at 37 °C using the high-resolution Oxygraph-2k (Oroboros, Innsbruck, Austria). Due to the low amount of oxygen required, this analysis was performed at oxygen concentrations between 200 (atmospheric pressure) and 60 nmol mL^-1^. The SUIT protocol used for this analysis was the same as the one utilized for determination of mitochondrial respiration in permeabilized kidney cortex, as described above. Data are presented as mitochondrial-specific respiration (pmol O_2_ s^-1^ mg^-1^ protein/CS activity, with CS activity determined in isolated mitochondria fractions).

### Determination of enzymatic activity

The enzymatic activity of Citrate synthase (CS) and Complex I (CI) were determined spectrophotometrically in mitochondria-enriched fractions (post-8000 g pellets) from human kidney cortices, as previously described ^24,25^, with minor modifications. Isolated mitochondrial fractions were diluted to a concentration of 2 mg/mL and sonicated with a 10 sec on, 10 sec off cycle at 90% amplitude at 4 °C for 6 min.

Due to the low mitochondrial yield and the good in-house reliability of the assay ^26^, CS activity was determined in duplicate on a microtiter plate by addition of 10 µL of sample to the following reaction mixture: 165 µL of 100 mM tris buffer (pH 8.3), 40 µL of 3 mM acetyl coenzyme A, 25 µL of 1 mM 5,5-dithiobis(2-nitrobenzoic acid) (DTNB) in tris buffer, and 10 µL of 10% Triton-X 100, kept at 25 °C. Endogenous CS activity was first determined in a spectrophotometer (CLARIOstar, BMG Labtech) after 30 s of linear agitation recording every 12 s for 3 min, at an absorbance of 412 nm at 25 °C. Maximal CS activity was then measured with the same parameters, immediately after the addition of 20 µL of 7.5 mM oxaloacetate. CS activity was reported as the difference between the two reads and expressed as mmol min^-1^ mg prot^-1^.

CI activity was determined in triplicate on a UV-transparent flat bottom microtiter plate by addition of the following reaction mixture: 150 μL of 2x sucrose/tris buffer (500 mM sucrose, 100 mM tris, 2 mM EDTA, pH to 7.4), 3 μL of 3 mM decylubiquinone (DB), 3 μL of 200 mM KCN, 15 μL of 1 mM NADH, 3 μL of 1 mM rotenone, 121 μL of MilliQ H_2_O (for a total of 295 μL), to 5 µL of sample, to determine the non CI-driven oxidation of NADH. In parallel, 295 μL of the same reaction mixture, where the 3 μL of 1 mM rotenone was replaced by 3 μL of EtOH, was added to three further wells containing 5 µL of sample, to determine the CI-driven oxidation of NADH, and immediately placed in the spectrophotometer (CLARIOstar, BMG Labtech), where both CI-driven and non CI-driven activities were simultaneously determined after 30 s of linear agitation recording every 8 s for 3 min, at an absorbance of 340 nm at 25 °C. Maximal CI activity was reported as the difference between CI- and non CI-driven activity (both expressed as mmol min^-1^ mg prot^-1^) and subsequently normalised by the corresponding value of CS activity determined above. USA).

### Statistical analysis

All values are reported as means ± SEM, unless otherwise specified. Outliers were first removed using the ROUT method set at Q = 1% ^27^. Differences between Ctrl and T2D were analysed by unpaired t-test with Welch’s correction when dataset was normally distributed (Shapiro-Wilk test *P* > 0.05), and by Mann-Whitney test when not normally distributed (Shapiro-Wilk test *P* < 0.05). Statistical details can be found in the associated figure legends. The level of statistical significance was set a priori at *P* < 0.05. GraphPad Prism (v. 9.1.1) for all statistical analyses.

## Results

To conduct a comprehensive metabolic analysis of the kidney from living human donors with normoglycemia and diabetes, freshly collected human kidneys were obtained from 50 live donors during tumour nephrectomy for renal cell carcinoma (Figure S1). Non-tumour parenchyma of the kidney cortex, as appropriately selected by an Anatomical Pathologist and confirmed to be clear of the tumour margin, was collected and immediately transported to the research laboratory (Figure 1A). Of the 50 live donors, 31 participants had normal glucose tolerance and were designated controls; 17 had T2D; one had type 1 diabetes and one had pre-diabetes defined by a fasting blood glucose of 6.2 mmol/l. The two latter participants were excluded from the study (Figure S1). Patients were matched for age, sex and kidney function (Table 1), as determined by estimated glomerular filtration rate (eGFR)^28^. As expected, BMI was increased in the diabetic subjects; importantly, eGFR was not different (Table 1).

**Figure 1.**
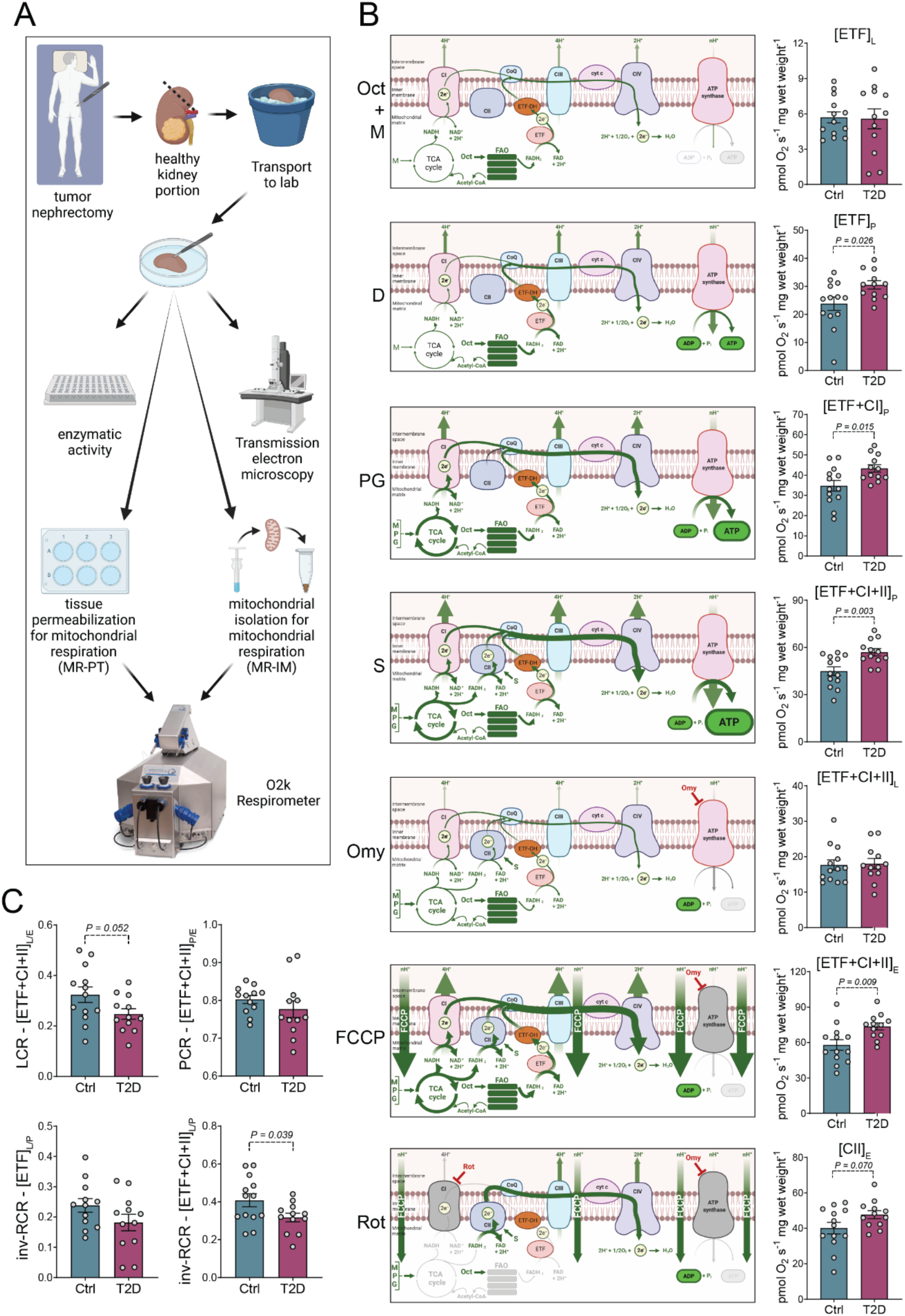
Diabetes-induced alterations in mitochondrial bioenergetics in the human kidney. (A) Study workflow; (B) mitochondrial respiration assessed in permeabilized human kidney cortex following addition of octanoylcarnitine (Oct), malate (M), ADP (D), pyruvate and glutamate (PG), succinate (S), cytochrome *c* (not shown), oligomycin (Omy), titration with carbonylcyanide 4-(trifluoromethoxy) phenylhydrazone (FCCP), rotenone (Rot), and antimycin a (not shown); (C) respiratory flux control ratio: leak control ratio (LCR), phosphorylation control ratio (PCR), and inverse respiratory control ratio (inv-RCR). ETF: electron transfer flavoprotein; L: leak respiration; CI: complex I; P OXPHOS respiration state; CII: complex II; E: electron transport system capacity; Ctrl: control patients; T2D: type 2 diabetic patients. Bars and dots represent mean and individual values, respectively; error bars represent SEM; n=13 (Ctrl) and 12 (T2D). Group differences were assessed by unpaired t-test with Welch’s correction when normally distributed or by Mann-Whitney test when non-normally distributed.

**Table 1.** Physiological and clinical characteristics of participants.

| Endpoint | Group | Age | Sex | BMI | eGFR |
| --- | --- | --- | --- | --- | --- |
| Total study | Ctrl | 61.1±12.6 | 18M - 13F | 27.7±6.8 | 87.4±13.6 |
|  | T2D | 62.5±7.4 | 9M - 8F | 32.9±7.9* | 89.9±9.8 |
| MR-PT | Ctrl | 64.8±9.9 | 7M - 6F | 27.8±7.5 | 82.0±11.9 |
|  | T2D | 61.2±7.9 | 7M - 5F | 32.1±8.7 | 89.8±11.1 |
| TEM | Ctrl | 60.8±12.6 | 10M - 8F | 27.2±6.3 | 88.8±12.6 |
|  | T2D | 61.2±7.6 | 8M - 6F | 32.0±8.1 | 90.1±10.8 |
| MR-IM | Ctrl | 59.6±16.5 | 6M - 4F | 28.0±7.6 | 94.8±13.0 |
|  | T2D | 62.4±5.8 | 4M - 6F | 33.2±9.1 | 90.6±9.2 |
| Enzyme activity | Ctrl | 62.4±12.3 | 17M - 11F | 28.2±6.8 | 86.9±13.6 |
|  | T2D | 61.8±7.6 | 9M - 6F | 31.0±6.0 | 90.1±10.5 |

\* $P < 0.05$ compared to control. BMI: body mass index; eGFR ( $\text{mL min}^{-1} 1.73 \text{ m}^2$ ): estimated glomerular filtration rate; Ctrl: healthy control participants; T2D: subjects with type 2 diabetes; TEM: transmission electron microscopy; MR: mitochondrial respiration; PT: permeabilized tissue; IM: isolated mitochondria.

### Diabetes is associated with an increased mitochondrial respiratory flux in permeabilized human kidney cortex

Measurement of oxidative phosphorylation (OXPHOS), involving the electron transport system (ETS) and complex V, by high-resolution respirometry enables detailed assessment of mitochondrial bioenergetics. Alterations in kidney mitochondrial bioenergetics have been widely reported in experimental diabetes and vary temporally, spatially and between models according to metabolic context, as recently reviewed ^29^. We first measured mitochondrial respiration in permeabilized kidney cortex. T2D was associated with increases in all three ATP-generating, coupled respiratory states measured. These included a 28% increase in OXPHOS capacity with electron input through electron-transferring flavoprotein (ETF), representative of fatty acid β-oxidation-linked respiration ([ETF]P; P=0.026); a 24% increase in OXPHOS capacity with convergent electron input through ETF and complex I (CI) ([ETF+CI]P; P=0.015); and a 27% increase in maximal coupled mitochondrial respiration with convergent electron input through ETF, CI and complex II (CII) ([ETF+CI+II]P; P=0.003) (Figure 1B). Maximal non-coupled ETS capacity ([ETF+CI+II]E) was also increased in T2D by 29% (P=0.009; Figure 1B), whereas non-coupled respiration supported by CII alone ([CII]E) was unchanged (P>0.05; Figure 1B). Neither of the two non-phosphorylating leak respiration states differed between groups ([ETF]L and [ETF+CI+II]L; both P>0.05; Figure 1B).

To further characterise respiratory control independently of absolute differences in mitochondrial abundance, we calculated flux control ratios (FCRs), which express individual respiratory states relative to maximal respiratory capacity within the same experiment. T2D was associated with a 22% reduction in the inverse respiratory control ratio (inv-RCR; P=0.039; Figure 1C). The leak control ratio (LCR), which reflects the proportion of maximal respiratory capacity attributable to non-phosphorylating respiration, was similarly reduced by 24% in T2D, although this did not reach statistical significance (P=0.052). These reductions were driven by increased OXPHOS and ETS capacity without a corresponding change in absolute leak respiration. Collectively, these findings demonstrate increased tissue-level mitochondrial respiratory flux in permeabilised human diabetic kidney cortex.

### Increased respiratory flux in permeabilised human kidney cortex is accounted for by greater mitochondrial volume density

The kidney has a high energetic requirement to support its physiological functions ^30^. Most of this energy is generated during OXPHOS in the mitochondria of proximal tubule epithelial cells (PTECs) ^31^, which are abundant in the kidney cortex ^30^.

Experimental models of diabetes demonstrate alterations in both mitochondrial bioenergetics and PTEC mitochondrial morphology ^32–36^. Mitochondrial structure and abundance are closely linked to cellular function and energy demand ^37,38^. We therefore investigated whether the increased respiratory flux observed in diabetic kidney cortex was associated with greater mitochondrial volume density (Mito_VD_). Mitochondrial morphology and abundance were assessed in fixed renal cortex by transmission electron microscopy (TEM), the gold-standard approach for quantifying mitochondrial content ^39^. T2D was associated with a 17% increase (*P*=0.002) in Mito_VD_ (Figure 2A and B). Further analysis demonstrated a 3% increase in mitochondrial roundness (*P*=0.002) and a trend towards a reduction in aspect ratio (-2%; *P*=0.088) (Figure 2C), consistent with mitochondrial fragmentation, as previously demonstrated in experimental models of diabetes ^32,34–36^. No differences were observed in the other mitochondrial shape descriptors examined (all *P*>0.05; Figure 2C & Figure S2).

**Figure 2.**
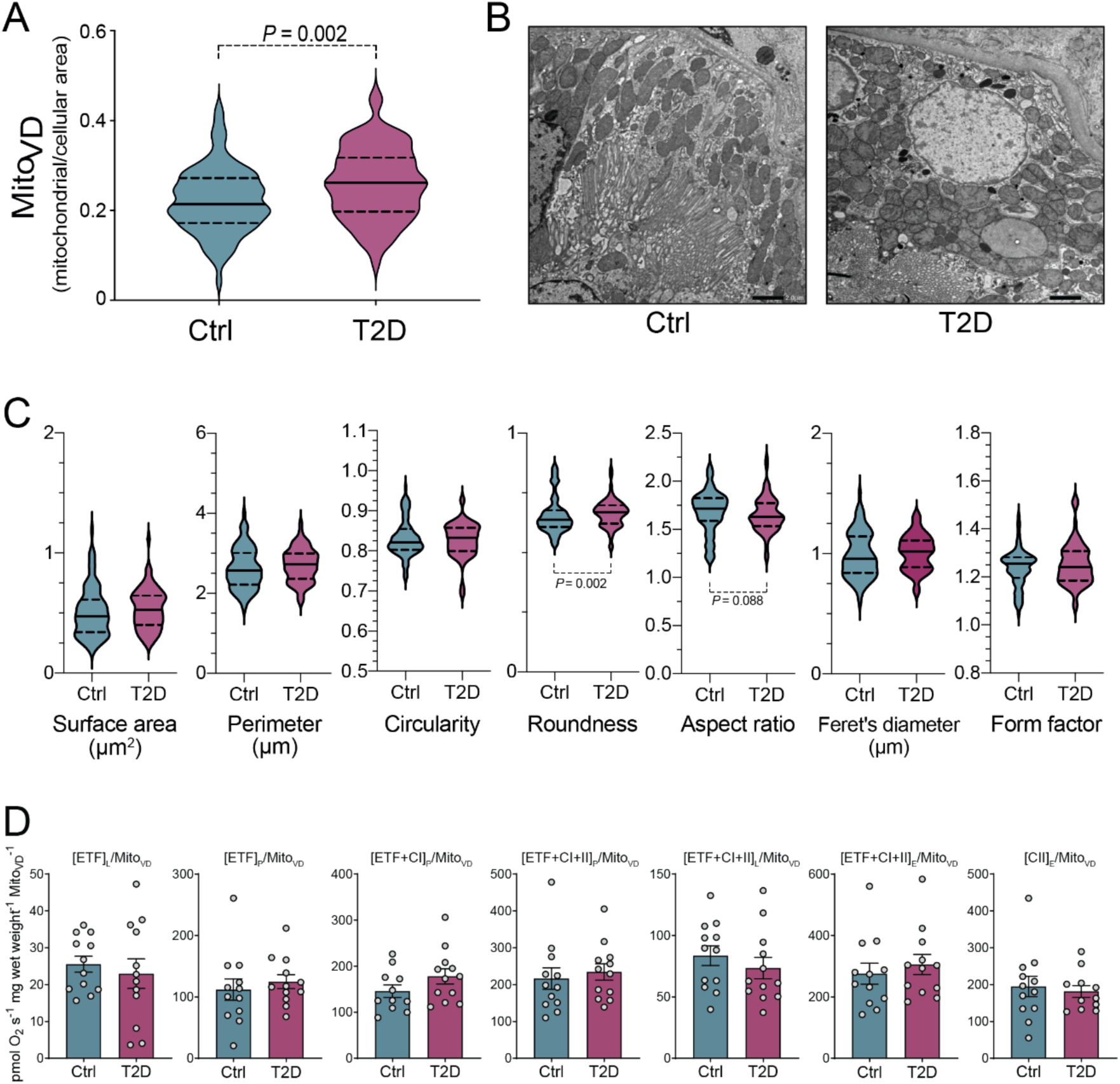
The diabetes-induced increase in mitochondrial respiration in human kidney cortex is mediated by increased mitochondrial volume density. Quantification of different parameters of mitochondrial morphology in Ctrl and T2D subjects from transmission electron microscopy (TEM) images. (A) Mitochondrial volume density (Mito_VD_); (B) representative electron micrographs of renal PTECs; x2,000 magnification; scale bar=2 µm; (C) mitochondrial shape descriptors; (D) mitochondrial-(mt-) specific respiration obtained by normalizing values of mitochondrial respiration (Figure 1B) by Mito_VD_. A, B and C: n=18 Ctrl and 14 T2D; 5 images/patient; solid lines represent the median; dashed lines represent the 25 and 75 percentile. D: n=12/group; bars and dots represent mean and individual values, respectively; error bars represent SEM. Group differences were assessed by unpaired t-test with Welch’s correction when normally distributed or by Mann-Whitney test when non-normally distributed.

To distinguish quantitative changes in mitochondrial abundance from differences in respiratory performance, tissue respiration values shown in Figure 1B were normalised to Mito_VD_ to derive mitochondria-specific respiratory flux (Figure 2D). The differences between T2D and control cortex were no longer evident following normalisation to Mito_VD_ (Figure 2D). These findings indicate that the increased respiratory flux observed in permeabilised diabetic kidney cortex was accounted for by an overall increase in mitochondrial volume density.

### Diabetes is associated with reduced intrinsic respiratory capacity in mitochondria isolated from human kidney cortex

To our knowledge, this is the first published study to directly characterise mitochondrial respiratory flux in fresh human diabetic kidney cortex. Because previous studies examining renal mitochondrial respiratory function in experimental diabetes have predominantly used isolated mitochondria ^29,32,40,41^, we next assessed respiration in mitochondria freshly isolated from the same cortical tissue.

To account for the increase in Mito_VD_ observed by TEM, respiration in isolated mitochondria was normalised to citrate synthase (CS) activity, a widely used marker of mitochondrial content ^39^, measured in the same preparation (Figure S3). Mitochondria-specific respiration was reduced in all three OXPHOS states measured, although only [ETF]P/CS and [ETF+CI+II]P/CS reached statistical significance (P=0.044 and P=0.030, respectively; Figure 3A). Both non-coupled ETS states were reduced by approximately 35% in T2D, with the reduction reaching statistical significance for [CII]E/CS (P=0.035) but not [ETF+CI+II]E/CS (P=0.056; Figure 3A). Neither leak respiration state differed between groups (both P>0.05; Figure 3A).

**Figure 3.**
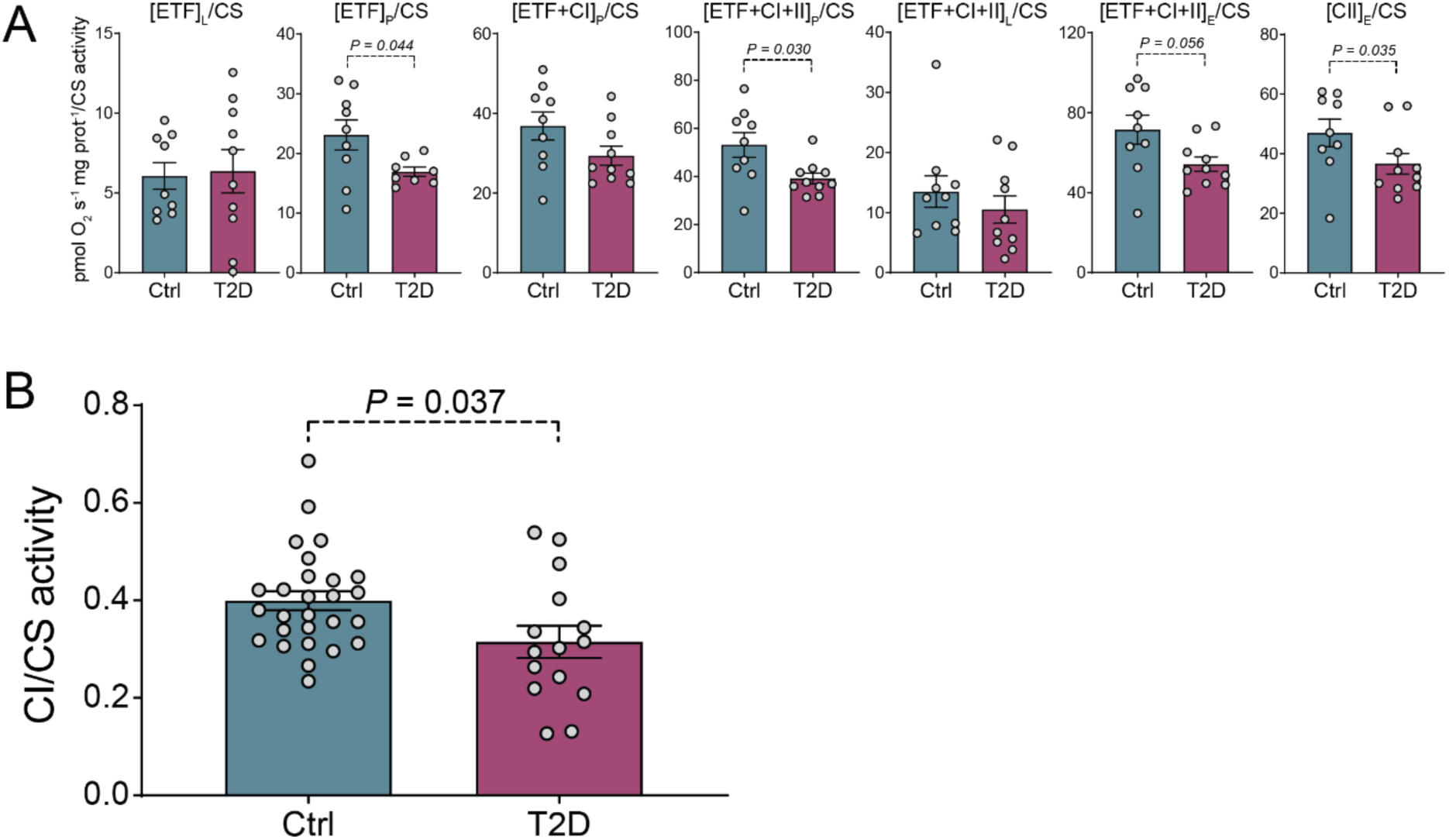
Diabetes induces a decrease in mitochondrial- (mt-) specific respiration and mitochondrial CI activity in mitochondria isolated from human kidney cortex. (A) Mitochondrial-specific respiration (obtained by normalizing mitochondrial respiration by citrate synthase [CS] activity) assessed in mitochondria isolated from human kidney cortex with the same protocol described in Figure 1B; (B) CI enzyme activity normalized by CS enzyme activity both determined in isolated mitochondria from human kidney cortex. Bars and dots represent mean and individual values, respectively; error bars represent SEM. A: n=10/group; B n=28 (Ctrl) and 15 (T2D). Group differences were assessed by unpaired t-test with Welch’s correction when normally distributed or by Mann-Whitney test when non-normally distributed.

Thus, despite increased respiratory flux at the tissue level, mitochondria isolated from diabetic kidney cortex demonstrated reduced intrinsic respiratory capacity when normalised to mitochondrial content. This discordance identifies a diabetic bioenergetic phenotype characterised by tissue-level hypermetabolism despite impaired mitochondria-specific function.

Experimental diabetes has also been associated with reduced activity of individual ETS complexes, particularly CI, in isolated kidney mitochondria ^40,42,43^. We therefore determined whether CI activity was similarly altered in mitochondria isolated from human kidney cortex. Following normalisation to CS activity to account for differences in mitochondrial abundance, CI activity was reduced in T2D (P=0.048; Figure 3B).

These findings provide direct evidence of intrinsic mitochondrial and CI dysfunction in the human diabetic kidney before measurable loss of kidney function.

## Discussion

The present study provides the first direct functional evidence that mitochondrial metabolism is reprogrammed in the human kidney during diabetes before measurable kidney dysfunction develops. Using freshly obtained kidney cortex from living donors, we demonstrate the unexpected coexistence of increased tissue-level mitochondrial respiratory flux with reduced intrinsic mitochondrial respiratory capacity and complex I activity. Together with increased mitochondrial density and fragmentation, these findings identify an early bioenergetic state characterised by tissue hypermetabolism despite intrinsic mitochondrial dysfunction. Rather than representing uniform mitochondrial failure, the diabetic kidney appears to undergo adaptive mitochondrial remodelling that sustains oxidative metabolism during the earliest stages of disease.

At first glance, increased respiratory flux in permeabilised cortex appears inconsistent with the reduced respiratory capacity observed in isolated mitochondria. However, these measurements interrogate distinct levels of biological organisation.

Permeabilised tissue reflects the integrated respiratory output of the mitochondrial network within intact proximal tubular cells, whereas isolated mitochondria quantify respiratory performance per mitochondrion. The increase in mitochondrial volume density, together with enhanced fragmentation, suggests expansion and remodelling of the mitochondrial network compensate for declining intrinsic mitochondrial efficiency. Consequently, total tissue respiration is maintained or even increased despite impaired respiratory performance of individual mitochondria.

Recent human studies have begun to identify early alterations in renal energy metabolism using imaging and single-cell approaches. In young adults with type 1 diabetes and preserved kidney function, ^11^C-acetate PET demonstrated attenuated cortical and medullary oxidative metabolism, accompanied by reduced proximal tubular oxidative phosphorylation and TCA-cycle transcripts and lower tubular TCA-cycle metabolites^44^. In contrast, single-cell profiling of renal biopsies from youth-onset type 2 diabetes identified activation of proximal tubular metabolic and mTORC1 pathways^45^, highlighting heterogeneity in the reported metabolic response of the human diabetic kidney. Our findings reconcile these apparently divergent observations by demonstrating that tissue-level respiratory output and intrinsic mitochondrial function represent distinct, and potentially opposing, features of early diabetic metabolic reprogramming. Whereas PET assesses integrated in vivo oxidative metabolism and transcriptomic analyses infer metabolic pathway activity, high-resolution respirometry directly quantifies mitochondrial respiratory function in viable tissue. The coexistence of increased cortical respiratory flux with reduced mitochondria-specific capacity therefore reveals a level of bioenergetic complexity that cannot be resolved by imaging or molecular profiling alone.

In the current study, the coexistence of increased tissue respiratory flux with impaired intrinsic mitochondrial function suggests that the early diabetic kidney undergoes compensatory mitochondrial remodelling rather than uniform bioenergetic failure. We propose that expansion of the mitochondrial network, evidenced by increased mitochondrial volume density, initially compensates for declining respiratory efficiency of individual mitochondria, thereby sustaining the exceptionally high energetic demands of the proximal tubule. This model is consistent with evidence that mitochondrial abundance adapts to tissue energy demand and contributes substantially to overall oxidative capacity^30^. In the diabetic kidney, increased mitochondrial abundance may therefore sustain tissue respiratory output despite reduced intrinsic respiratory performance. This adaptive state may preserve energy production early in diabetes but could become increasingly difficult to maintain as mitochondrial dysfunction progresses.

Increased mitochondrial volume density was accompanied by greater mitochondrial roundness, consistent with enhanced mitochondrial fragmentation. Mitochondrial fragmentation has generally been considered a pathological feature of diabetic kidney disease and has been linked to impaired bioenergetics, oxidative stress and tubular injury in experimental models^13,46^. However, fragmentation is also an integral component of mitochondrial quality control, facilitating mitochondrial turnover, redistribution and biogenesis under conditions of increased energetic demand^41^. Our findings suggest that, in the absence of measurable kidney dysfunction, increased mitochondrial fragmentation may represent an adaptive component of mitochondrial network remodelling rather than simply a marker of injury. Whether this adaptive response ultimately becomes maladaptive as diabetes progresses remains to be determined.

Reduced complex I activity provides direct biochemical evidence that intrinsic mitochondrial dysfunction is already present in the diabetic kidney before measurable loss of renal function. Complex I is the major entry point for NADH-derived electrons into the electron transport system, and reduced complex I activity and complex I-supported respiration have been reported in experimental diabetic kidney disease^13,47^. Consistent with a causal role, our previous Ndufs6 study demonstrated that targeted disruption of a complex I subunit directly impairs renal mitochondrial function and modifies the kidney injury phenotype^48^. Our current findings demonstrate that this defect is also present in human diabetic kidney cortex and is accompanied by reduced mitochondria-specific respiratory capacity. Importantly, these abnormalities were only apparent after accounting for differences in mitochondrial abundance, highlighting the importance of distinguishing intrinsic mitochondrial performance from overall tissue respiratory output.

Our findings also provide a framework for interpreting the apparently conflicting literature describing renal mitochondrial metabolism in diabetes. Experimental studies performed in isolated mitochondria have consistently concluded that mitochondrial respiration is impaired, whereas recent human imaging and transcriptomic studies have variably reported reduced oxidative metabolism or activation of metabolic pathways^49^. Rather than representing conflicting biological phenomena, these observations likely reflect different levels of biological organisation. Isolated mitochondria quantify respiratory capacity per mitochondrion, whereas tissue respiration reflects the integrated output of the entire mitochondrial network within its cellular environment. Consequently, increased tissue respiratory flux and impaired intrinsic mitochondrial function are not mutually exclusive but instead define complementary features of early diabetic metabolic reprogramming.

These findings have important implications for therapeutic strategies targeting mitochondrial function in diabetic kidney disease. Mitochondrial dysfunction has generally been viewed as a progressive loss of oxidative capacity requiring restoration of mitochondrial respiration^30,33^. Our data suggest a more nuanced model in which oxidative metabolism is initially maintained through expansion and remodelling of the mitochondrial network despite declining intrinsic respiratory efficiency. Therapeutic approaches aimed solely at increasing mitochondrial respiration may therefore overlook an early compensatory state in which tissue oxidative metabolism is already elevated. Instead, preserving mitochondrial quality, limiting metabolic stress and maintaining efficient respiratory function may represent more appropriate strategies during the earliest stages of DKD.

The principal strengths of this study include direct functional assessment of freshly obtained human kidney cortex from living donors with preserved kidney function, thereby avoiding many of the limitations associated with archived tissue or samples obtained from deceased donors. The combination of high-resolution respirometry in permeabilised tissue, isolated mitochondrial bioenergetics, quantitative transmission electron microscopy and respiratory chain enzyme activity enabled interrogation of mitochondrial function across multiple biological scales within the same human tissue samples. This integrated approach provides a comprehensive functional characterisation of mitochondrial metabolism that has not previously been possible in the human diabetic kidney.

Several limitations should be acknowledged. The cross-sectional design precludes assessment of temporal changes in mitochondrial adaptation or determination of whether the compensatory state identified here ultimately progresses to overt mitochondrial failure during diabetic kidney disease. Tissue was obtained from tumour nephrectomy specimens, although all samples were collected from histopathologically normal cortex remote from the tumour margin. The study predominantly included participants with type 2 diabetes and preserved kidney function, and therefore the findings may not be generalisable to advanced diabetic kidney disease or type 1 diabetes. Finally, although our data identify a coherent bioenergetic phenotype, the molecular mechanisms underpinning this adaptive mitochondrial remodelling require further investigation.

## Conclusion

Collectively, these findings redefine the early mitochondrial phenotype of the human diabetic kidney. Rather than exhibiting uniform bioenergetic failure, the kidney enters a previously unrecognised adaptive state characterised by increased tissue respiratory flux despite impaired intrinsic mitochondrial function. We propose that expansion and remodelling of the mitochondrial network initially compensate for declining mitochondrial efficiency to sustain the extraordinary energetic demands of the kidney. Failure of this adaptive state may represent a critical transition toward diabetic kidney disease.

## Author Contributions

SGW, CML, PR, JG, and MTC developed the workflow for the kidney biospecimen procurement at Alfred Health. JG, CC and NC performed nephrectomy. EIE supervised the kidney biospecimen procurement in conjunction with the Victorian Cancer Biobank. CG, AL, VTB, GR conducted experiments. CG and AL performed data analysis. CG and MTC wrote the manuscript. RJM, MEC, CML and SGW critically reviewed the manuscript. MTC conceptualized and supervised the studies and procured the funding.

## Acknowledgments

We thank the following people for their input Dr Simon Crawford, Dr David Stroud, Dr Gavin Higgins, Dr Runa Lindblom, Mrs Maryann Arnstein, Associate Professor Ian Trounce, Ms Lingyun Kong and Professor Rowan Walker. We thank the Urology Perioperative Coordinator Ms Anna Paton and nursing staff. We acknowledge the Ramaciotti Centre for Cryo-Electron Microscopy, Monash Micro Imaging and the Monash University Histology Service. We thank all staff at the Victorian Cancer Biobank and associated Urological surgeons and we are indebted to the participants who donated their kidney specimens to this research study. The Victorian Cancer Biobank through the Cancer Council Victoria as lead agency is supported by the Victorian Government through the Victorian Cancer Agency, a business unit of the Department of Health. This work was supported by the following agencies: JDRF Australia Type 1 Diabetes Clinical Research Network and the National Health and Medical Research Council of Australia (NHMRC, GNT2030559). MTC was the recipient of a Career Development Award from JDRF Australia (4-CDA-2018-613-M-B) through the Type 1 Diabetes Clinical Research Network, a special initiative of the Australian Research Council. MEC was a recipient of a NHMRC Investigator Grant. Figures were created with BioRender.com.

## SUPPLEMENTAL MATERIALS

### SUPPLEMENTARY FIGURES

**Figure S1.**
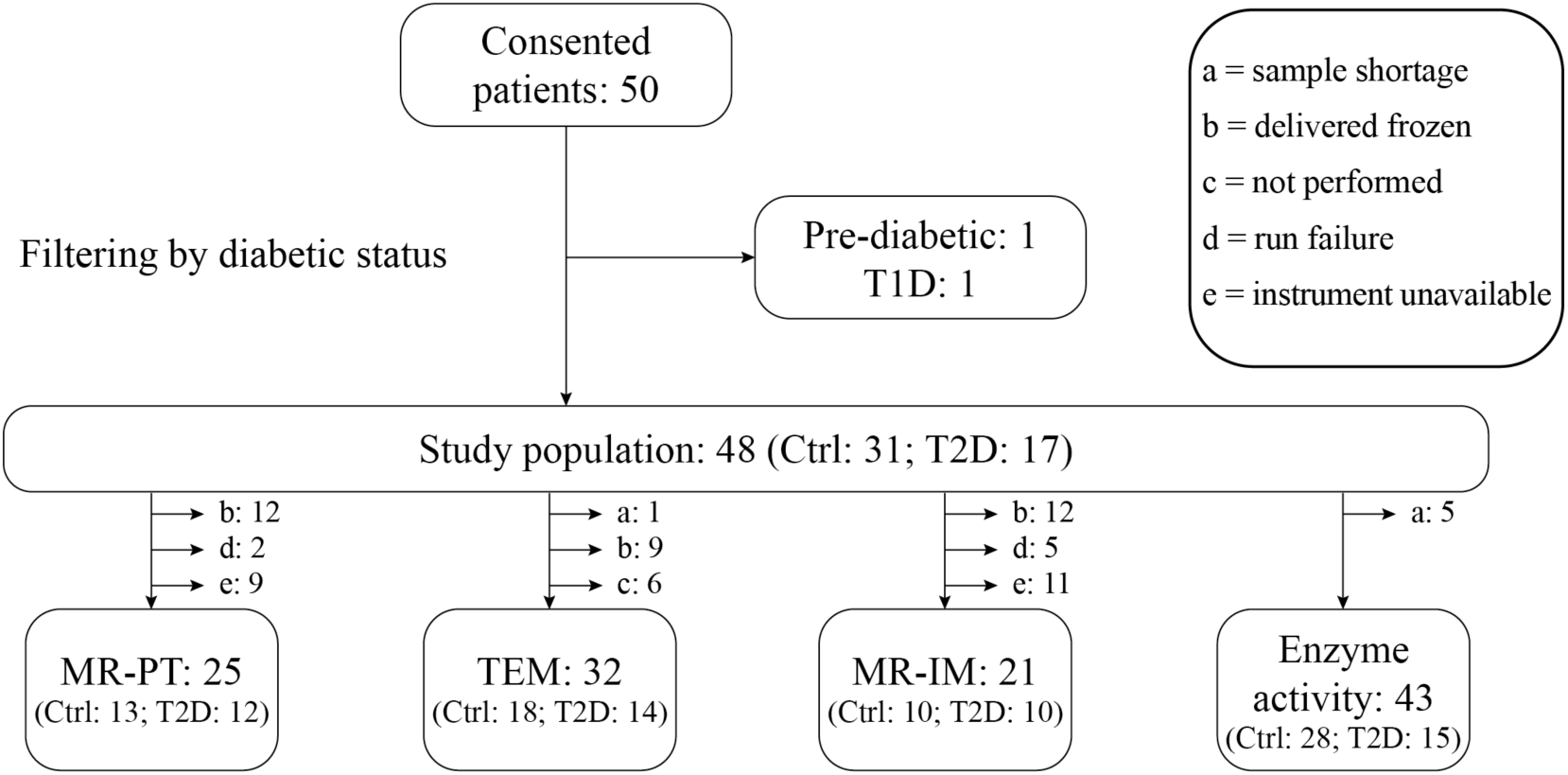
Study flow chart and analysis breakdown. eGFR: estimated glomerular filtration rate (expressed in mL min^-1^ 1.73 m^2^); T1D: type 1 diabetes; Ctrl: control participants; T2D: subjects with type 2 diabetes; TEM: transmission electron microscopy; MR: mitochondrial respiration; PT: permeabilized tissue; IM: isolated mitochondria.

**Figure S2.**
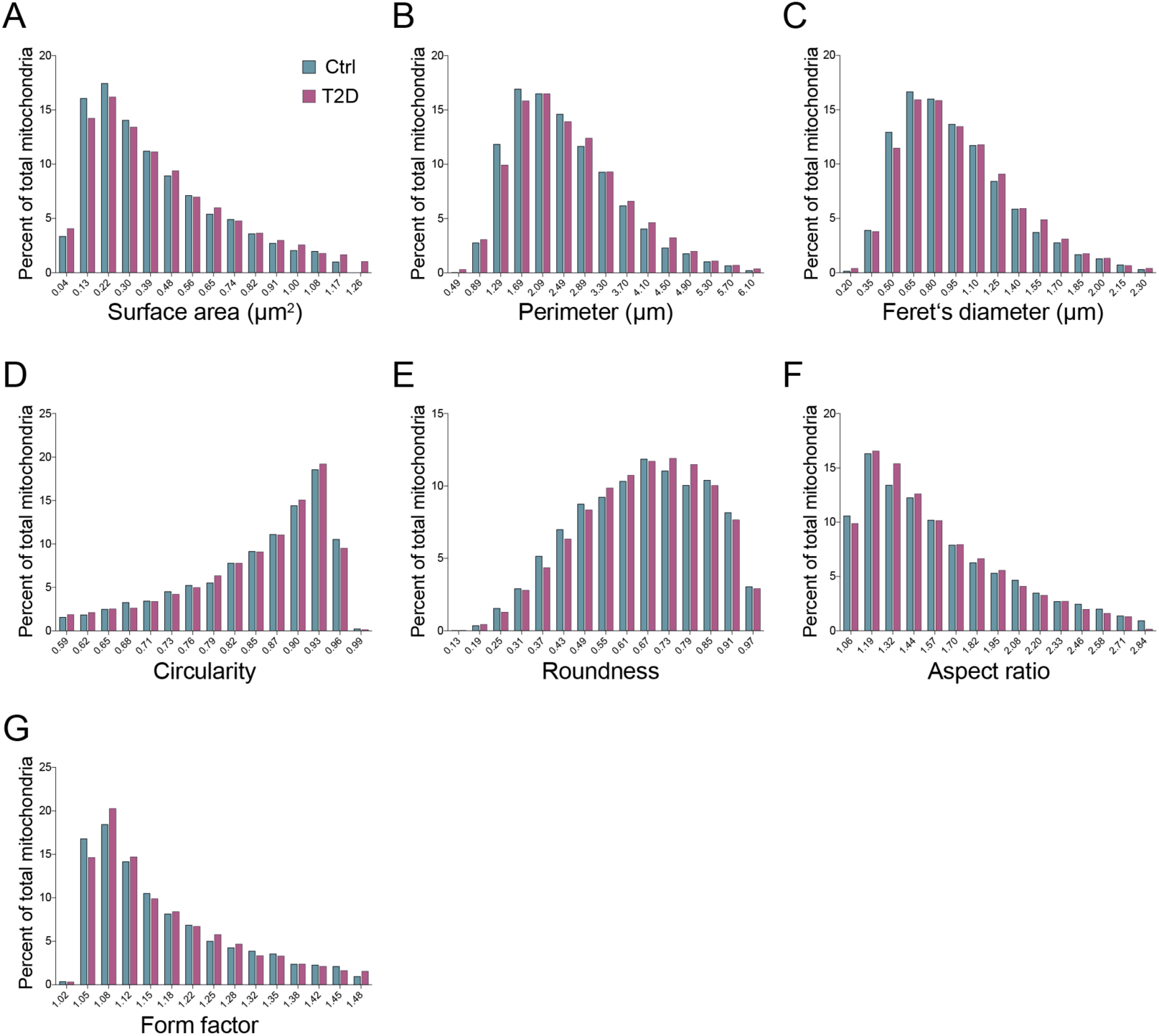
Frequency distribution plots for the seven shape descriptors presented in Figure 2C. Frequency distribution plots of morphological parameters from human proximal tubule epithelial cells mitochondria. Frequency distribution (% of total mitochondria) was plotted for (A) surface area, (B), perimeter, (C) Feret’s diameter, (D) circularity, (E) roundness, (F) aspect ratio, and (G) form factor, with 15 bins of equal sizes; n = 10,343 (Ctrl) and 8,470 (T2D) mitochondria.

**Figure S3.**
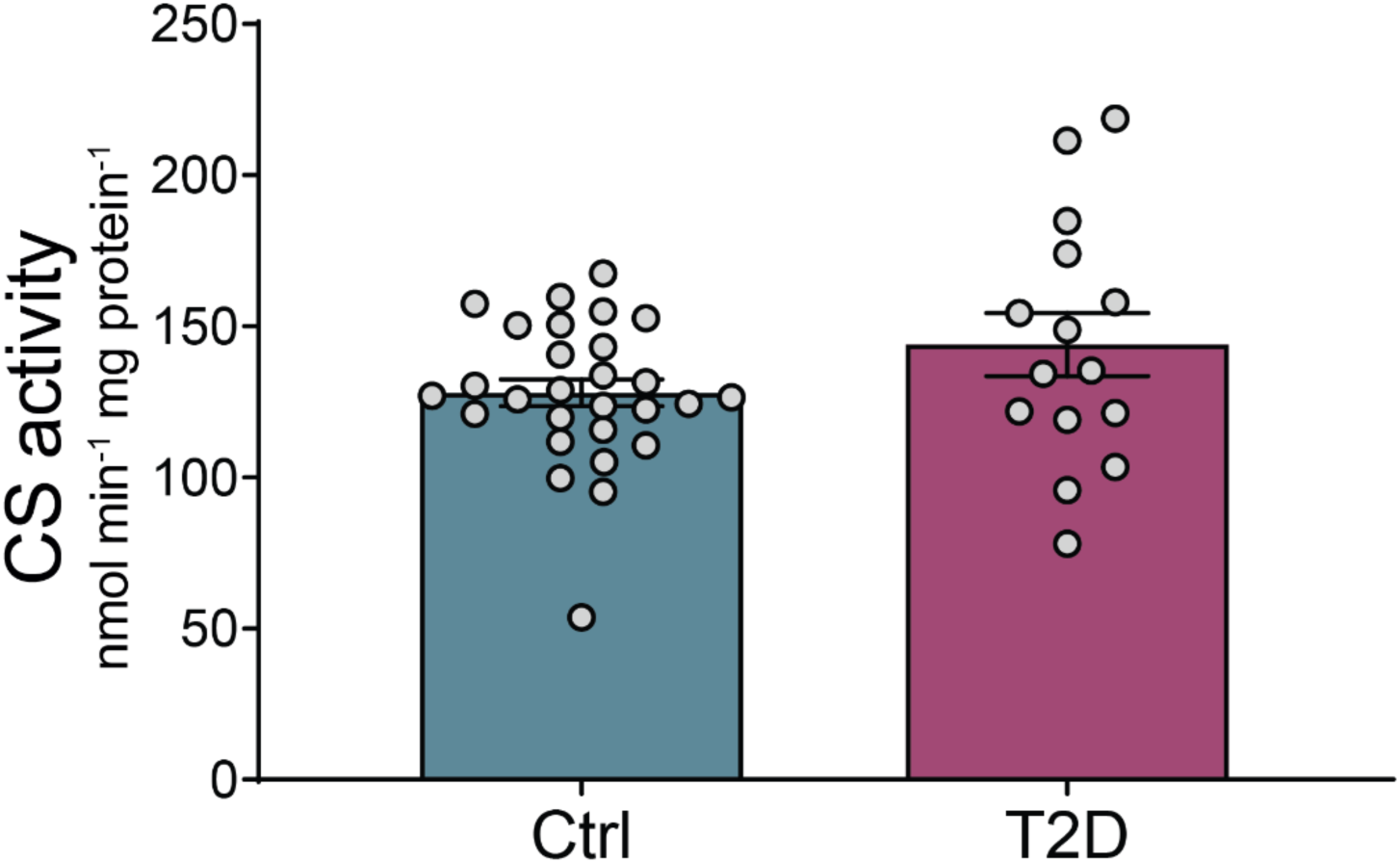
Citrate synthase (CS) activity in mitochondria isolated from human kidney cortex. Citrate synthase (CS) enzymatic activity assessed in mitochondrial-enriched fractions from human kidney cortex. Ctrl: control; T2D: type 2 diabetes. Bars and dots represent mean and individual values, respectively; error bars represent SEM; n=28 (Ctrl) and 15 (T2D). Group differences were assessed by unpaired t-test with Welch’s correction when normally distributed or by Mann-Whitney test when non-normally distributed. Significance set at *P*<0.05.

## References

1. International Diabetes Federation Diabetes Atlas 10th Edition. (2021).

2. Koye, D.N., Magliano, D.J., Nelson, R.G. & Pavkov, M.E. The Global Epidemiology of Diabetes and Kidney Disease. Adv Chronic Kidney Dis 25, 121–132 (2018).

3. Perkovic, V., et al. Canagliflozin and Renal Outcomes in Type 2 Diabetes and Nephropathy. N Engl J Med (2019).

4. Heerspink, H.J.L., et al. Dapagliflozin in Patients with Chronic Kidney Disease. N Engl J Med 383, 1436–1446 (2020).

5. Soltoff, S.P. ATP and the regulation of renal cell function. Annual review of physiology 48, 9–31 (1986).

6. Clark, A.J. & Parikh, S.M. Targeting energy pathways in kidney disease: the roles of sirtuins, AMPK, and PGC1alpha. Kidney Int 99, 828–840 (2021).

7. Kang, H.M., et al. Defective fatty acid oxidation in renal tubular epithelial cells has a key role in kidney fibrosis development. Nature medicine 21, 37–46 (2015).

8. Sivitz, W.I. & Yorek, M.A. Mitochondrial dysfunction in diabetes: from molecular mechanisms to functional significance. Antioxidants & redox signaling 12, 537–577 (2010).

9. Daehn, I., et al. Endothelial mitochondrial oxidative stress determines podocyte depletion in segmental glomerulosclerosis. J Clin Invest 124, 1608–1621 (2014).

10. Che, R., Yuan, Y., Huang, S. & Zhang, A. Mitochondrial dysfunction in the pathophysiology of renal diseases. American journal of physiology. Renal physiology 306, F367–378 (2014).

11. Hall, A.M. & Unwin, R.J. The not so ‘mighty chondrion’: emergence of renal diseases due to mitochondrial dysfunction. Nephron Physiol 105, p1–10 (2007).

12. Coughlan, M.T., et al. Deficiency in Apoptosis-Inducing Factor Recapitulates Chronic Kidney Disease via Aberrant Mitochondrial Homeostasis. Diabetes 65, 1085–1098 (2016).

13. Coughlan, M.T., et al. Mapping time-course mitochondrial adaptations in the kidney in experimental diabetes. Clin Sci (Lond) 130, 711–720 (2016).

14. Coughlan, M.T., et al. Combination therapy with the advanced glycation end product cross-link breaker, alagebrium, and angiotensin converting enzyme inhibitors in diabetes: synergy or redundancy? Endocrinology 148, 886–895 (2007).

15. Coughlan, M.T., et al. RAGE-induced cytosolic ROS promote mitochondrial superoxide generation in diabetes. J Am Soc Nephrol 20, 742–752 (2009).

16. Tan, A.L., et al. Disparate effects on renal and oxidative parameters following RAGE deletion, AGE accumulation inhibition, or dietary AGE control in experimental diabetic nephropathy. American journal of physiology. Renal physiology 298, F763–770 (2010).

17. Tan, S.M., et al. Complement C5a Induces Renal Injury in Diabetic Kidney Disease by Disrupting Mitochondrial Metabolic Agility. Diabetes 69, 83–98 (2020).

18. Sharma, K., et al. Metabolomics reveals signature of mitochondrial dysfunction in diabetic kidney disease. Journal of the American Society of Nephrology 24, 1901–1912 (2013).

19. Gordin, D., et al. Characterization of Glycolytic Enzymes and Pyruvate Kinase M2 in Type 1 and 2 Diabetic Nephropathy. Diabetes Care 42, 1263–1273 (2019).

20. Qi, W., et al. Pyruvate kinase M2 activation may protect against the progression of diabetic glomerular pathology and mitochondrial dysfunction. Nature medicine 23, 753–762 (2017).

21. Pena, M.J., et al. The effects of atrasentan on urinary metabolites in patients with type 2 diabetes and nephropathy. Diabetes Obes Metab 19, 749–753 (2017).

22. Picard, M., White, K. & Turnbull, D.M. Mitochondrial morphology, topology, and membrane interactions in skeletal muscle: A quantitative three-dimensional electron microscopy study. J. Appl. Physiol. 114, 161–171 (2013).

23. Granata, C., Oliveira, R.S.F., Little, J.P., Renner, K. & Bishop, D.J. Mitochondrial adaptations to high-volume exercise training are rapidly reversed after a reduction in training volume in human skeletal muscle. FASEB J. 30, 3413–3423 (2016).

24. Granata, C., Oliveira, R.S.F., Little, J.P., Renner, K. & Bishop, D.J. Training intensity modulates changes in PGC-1α and p53 protein content and mitochondrial respiration, but not markers of mitochondrial content in human skeletal muscle. FASEB J. 30, 959–970 (2016).

25. Van Bergen, N.J., Blake, R.E., Crowston, J.G. & Trounce, I.A. Oxidative phosphorylation measurement in cell lines and tissues. Mitochondrion 15, 24–33 (2014).

26. Kuang, J., et al. Methodological considerations when assessing mitochondrial respiration and biomarkers for mitochondrial content in human skeletal muscle. bioRxiv (2021).

27. Motulsky, H.J. & Brown, R.E. Detecting outliers when fitting data with nonlinear regression–a new method based on robust nonlinear regression and the false discovery rate. BMC Bioinformatics 7, 123 (2006).

28. Levey, A.S., et al. A new equation to estimate glomerular filtration rate. Ann Intern Med 150, 604–612 (2009).

29. Mise, K., Galvan, D.L. & Danesh, F.R. Shaping Up Mitochondria in Diabetic Nephropathy. Kidne*y360* 1, 982–992 (2020).

30. Bhargava, P. & Schnellmann, R.G. Mitochondrial energetics in the kidney. Nat Rev Nephrol 13, 629–646 (2017).

31. Soltoff, S.P. ATP and the regulation of renal cell function. Annu. Rev. Physiol. 48, 9–31 (1986).

32. Coughlan, M.T., et al. Mapping time-course mitochondrial adaptations in the kidney in experimental diabetes. Clinical science 130, 711–720 (2016).

33. Forbes, J.M. & Thorburn, D.R. Mitochondrial dysfunction in diabetic kidney disease. Nature Reviews Nephrology 14, 291 (2018).

34. Galloway, C.A., et al. Transgenic control of mitochondrial fission induces mitochondrial uncoupling and relieves diabetic oxidative stress. Diabetes 61, 2093–2104 (2012).

35. Wang, W., et al. Mitochondrial fission triggered by hyperglycemia is mediated by ROCK1 activation in podocytes and endothelial cells. Cell metabolism 15, 186–200 (2012).

36. Zhan, M., Usman, I.M., Sun, L. & Kanwar, Y.S. Disruption of renal tubular mitochondrial quality control by Myo-inositol oxygenase in diabetic kidney disease. Journal of the American Society of Nephrology 26, 1304–1321 (2015).

37. Galvan, D.L., Green, N.H. & Danesh, F.R. The hallmarks of mitochondrial dysfunction in chronic kidney disease. Kidney international 92, 1051–1057 (2017).

38. Ryan, M.T. & Hoogenraad, N.J. Mitochondrial-nuclear communications. Annu. Rev. Biochem. 76, 701–722 (2007).

39. Larsen, S., et al. Biomarkers of mitochondrial content in skeletal muscle of healthy young human subjects. Journal of Physiology 590, 3349–3360 (2012).

40. Sharma, K. Mitochondrial dysfunction in the diabetic kidney. in Advances in Experimental Medicine and Biology, Vol. 982 553–562 (2017).

41. Higgins, G.C. & Coughlan, M.T. Mitochondrial dysfunction and mitophagy: the beginning and end to diabetic nephropathy? Br J Pharmacol 171, 1917–1942 (2014).

42. Coughlan, M.T., et al. RAGE-induced cytosolic ROS promote mitochondrial superoxide generation in diabetes. Journal of the American Society of Nephrology 20, 742–752 (2009).

43. Dugan, L.L., et al. AMPK dysregulation promotes diabetes-related reduction of superoxide and mitochondrial function. The Journal of clinical investigation 123(2013).

44. Choi, Y.J., et al. Attenuated kidney oxidative metabolism in young adults with type 1 diabetes. J Clin Invest 134(2024).

45. Schaub, J.A., et al. SGLT2 inhibitors mitigate kidney tubular metabolic and mTORC1 perturbations in youth-onset type 2 diabetes. J Clin Invest 133(2023).

46. Lee, Y.H., et al. Empagliflozin attenuates diabetic tubulopathy by improving mitochondrial fragmentation and autophagy. American journal of physiology. Renal physiology 317, F767–F780 (2019).

47. Sourris, K.C., et al. Ubiquinone (coenzyme Q10) prevents renal mitochondrial dysfunction in an experimental model of type 2 diabetes. Free Radic Biol Med 52, 716–723 (2012).

48. Forbes, J.M., et al. Deficiency in mitochondrial complex I activity due to Ndufs6 gene trap insertion induces renal disease. Antioxidants & redox signaling 19, 331–343 (2013).

49. Pinzon-Cortes, J.A., et al. Mitochondrial Alterations and CKD. Am J Kidney Dis (2026).

